# Ageing affects DNA methylation drift and transcriptional cell-to-cell variability in muscle stem cells

**DOI:** 10.1101/500900

**Authors:** Irene Hernando-Herraez, Brendan Evano, Thomas Stubbs, Pierre-Henri Commere, Stephen Clark, Simon Andrews, Shahragim Tajbakhsh, Wolf Reik

**Author notes:** Equal contributions (Irene Hernando-Herraez, Brendan Evano, Thomas Stubbs). Equal contributions (Shahragim Tajbakhsh, Wolf Reik).

## Abstract

Age-related tissue alterations have been associated with a decline in stem cell number and function^1^. Although increased cell-to-cell variability in transcription or epigenetic marks has been proposed to be a major hallmark of ageing^2–5^, little is known about the molecular diversity of stem cells during ageing. Here, by combined single-cell transcriptome and DNA methylome profiling in mouse muscle stem cells, we show a striking global increase of uncoordinated transcriptional heterogeneity together with context-dependent alterations of DNA methylation with age. Importantly, promoters with increased methylation heterogeneity are associated with increased transcriptional heterogeneity of the genes they drive. Notably, old cells that change the most with age reveal alterations in the transcription of genes regulating cell-niche interactions. These results indicate that epigenetic drift, by accumulation of stochastic DNA methylation changes in promoters, is a substantial driver of the degradation of coherent transcriptional networks with consequent stem cell functional decline during ageing.

---

Epigenetic alterations have been proposed to be a major cause of age-related decline in tissue function^6^. Changes in DNA methylation are well correlated with ageing and methylation of specific loci has been used as age biomarker in a large number of tissues^6,7^. However, age-related methylation changes are poorly correlated with transcriptional variation, presumably because the changes are generally small and may not occur homogeneously in all cells^7^, a phenomenon also known as epigenetic drift. Although epigenetic drift has long been hypothesised to be an important hallmark of ageing^8^, this proposal has been challenging to test because of technical constraints. However, powerful combined single cell methods^9,10^ are now available, and epigenetic changes during ageing together with their functional consequences can now be read out in single cells^11^.

Degenerative changes in tissue-specific stem cells have been proposed to be a major cause of age-related decline in tissue function^12^. While several reports indicate a loss of clonal diversity during early life stages^13–15^ little is known about how cell-to-cell variability at the molecular level is involved in stem cell ageing. Here, we performed parallel single-cell DNA methylation and transcriptome sequencing (scM&T-seq) on the same cell^10^ to investigate how ageing affects transcriptional and epigenetic heterogeneity of tissue-specific stem cells, using mouse muscle stem cells as a model. Muscle satellite (stem) cells express the transcription factor *Pax7*^16^ and are largely quiescent in adult muscles. They activate upon injury to differentiate and fuse to form new fibers, or self-renew to reconstitute the stem cell pool^16^. Age-associated muscle defects have been attributed to a decrease in stem cell number together with impaired regenerative potential^17^. In addition, clonal lineage-tracing of mouse satellite cells showed that population diversity is unaltered during homeostatic ageing^18^.

Satellite cells with high expression of *Pax7* were shown to be in a deep quiescent state^19,20^. To investigate the molecular effects of ageing in a defined population that is less poised to enter the cell cycle, we isolated single satellite cells by fluorescence-activated cell sorting (FACS) from young (2 months) and old (24 months) *Tg:Pax7-nGFP* mice^21^ and selected those with high levels of GFP, to which we applied scM&T-seq (Fig. 1A).

**Fig. 1.**
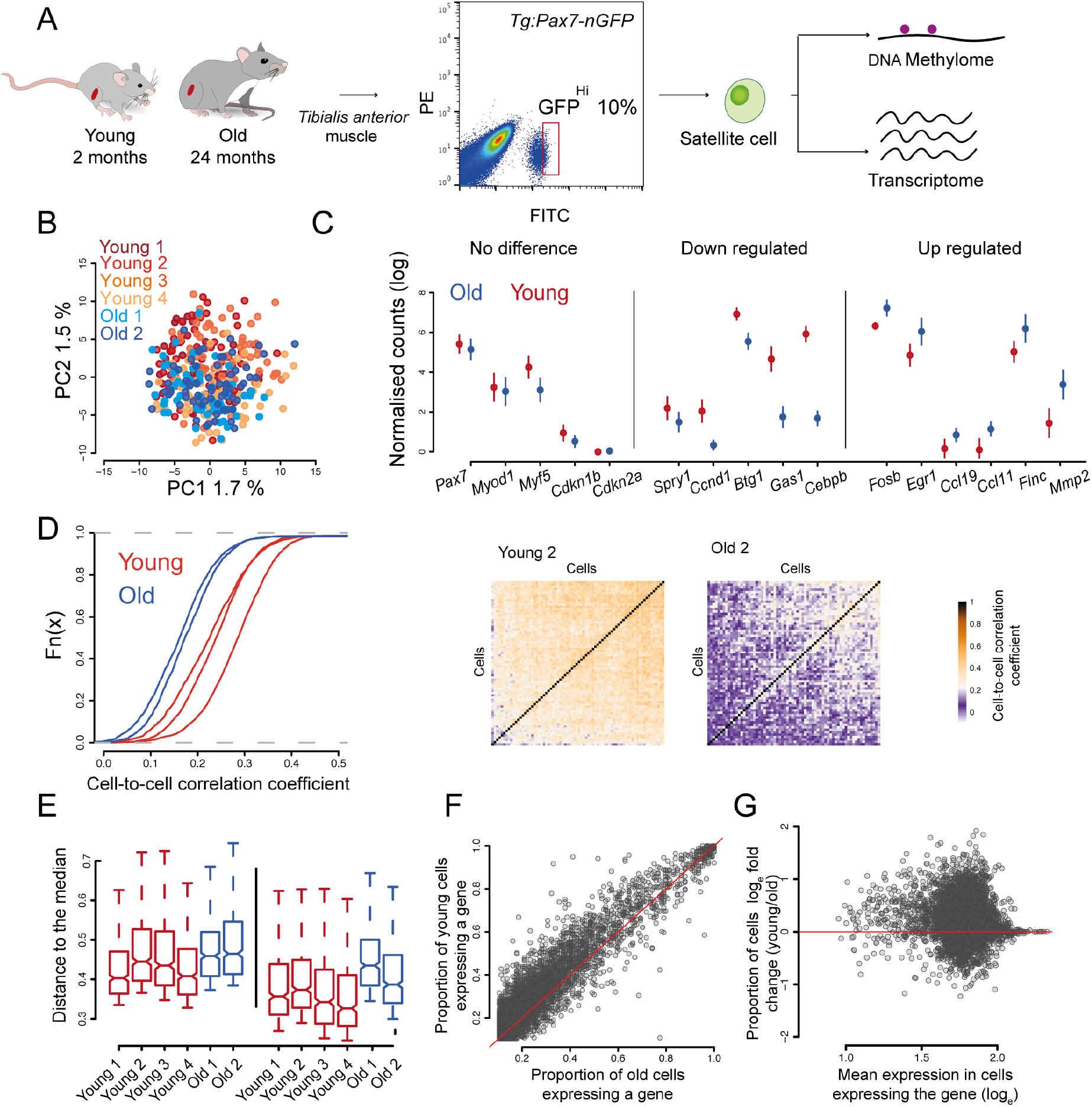
Aged satellite cells have increased cell-to-cell transcriptional variability. (A) Experimental scheme. Single cells were isolated from *Tg:Pax7-nGFP* young and old mice and subjected to parallel single-cell methylation and RNA sequencing. (B) PCA of a total of 377 cells from young (n=4) and old (n=2) individuals. (C) Selected markers and differentially expressed genes between young and old cells (mean ± standard error). (D) Cumulative distribution of cell-to-cell Spearman correlation values per individual (left) showing that transcriptional heterogeneity dramatically increases with age. Heatmap showing cell-to-cell Spearman correlation values from a young and an old mouse (right). (E) Distance to the median of the top 500 most variable genes among all genes (left) and of the top 500 most variable genes among the 5,127 common genes expressed in the six individuals (right). (F) Frequency of gene expression in young and old cells. (G) Independence between frequency of gene expression differences and gene expression level.

After quality control and filtering, a total of 377 transcriptomes were analysed. Young and old cells from different individuals clustered together, respectively, indicating no global differences with age and absence of sequencing-related batch effects (Fig. 1B). Furthermore, we did not observe significant differences in the levels of *Pax7*, the myogenic factors *Myod* and *Myf5* and the cell cycle inhibitor *Cdkn1b*, nor of senescent markers such as *Cdkn2a*, suggesting that some molecular signatures are conserved between the analysed cell populations (Fig. 1C). Nevertheless, 940 genes were differentially expressed between young and old individuals (SCDE, FDR *P* < 0.05, Table S1). *Spry1*, which is a key factor for maintaining quiescence^22^, and the cell cycle regulators *Ccnd1, Btg1* and *Gas1* were down-regulated, while ageing markers such as the chemokine genes *Ccl11* and *Ccl19* were up-regulated^20^ (Fig. 1C). Furthermore, we uncovered genes not previously reported to change in expression with age, such as the early activation markers *Fosb* and *Egr1*^23^ and the metalloproteinase *Mmp2* (Fig. 1C).

To investigate if ageing affects transcriptional heterogeneity of the stem cell pool, we calculated pairwise correlation coefficients between cells within each individual (see Methods) and observed that old individuals showed consistently lower correlation (1.3 mean-fold decrease, Mann-Whitney-Wilcoxon test; *P* < 2.2e-16, Fig. 1D), indicating a remarkably lower degree of similarity between cells and no obvious population substructure. We also computed an expression-level normalised measure of gene expression heterogeneity (named distance to the median)^24^, which proved to be higher in old individuals (Mann-Whitney-Wilcoxon test; *P* < 2.2e-16, Fig. 1E) revealing a striking global increase of uncoordinated transcriptional variability with age. Strikingly, the proportion of cells expressing a given gene (frequency of gene expression) was reduced with age (Mann-Whitney-Wilcoxon test; *P* < 2.2e-16, Fig. 1F), even in genes that did not significantly change mean expression levels (SCDE, FDR *P* > 0.05, Fig. 1F). Importantly, we observed that this was independent of gene expression levels and not restricted to lowly expressed genes suggesting that this global feature is unrelated to technical effects (Fig. 1G).

Genes that displayed increased expression variability with age (expression frequency difference > 15%) include several collagen genes (*Col4a2, Col5a3, Col4a1*) and other extracellular matrix-related genes such as *Dag1, Sparc, Cdh15* or *Itgb1* (Fig. 2A). Interestingly, satellite cells without *Itgb1* (β1-integrin) cannot maintain quiescence and its experimental activation improves ageing-related decline in muscle regeneration^25^. Similarly, reduction of N-cadherin and M-cadherin (*Cdh15*) leads to a break of quiescence of satellite cells^26^. Notably, none of the above-mentioned genes were shown to change in expression level during the isolation procedure of satellite cells^27^.

**Fig. 2.**
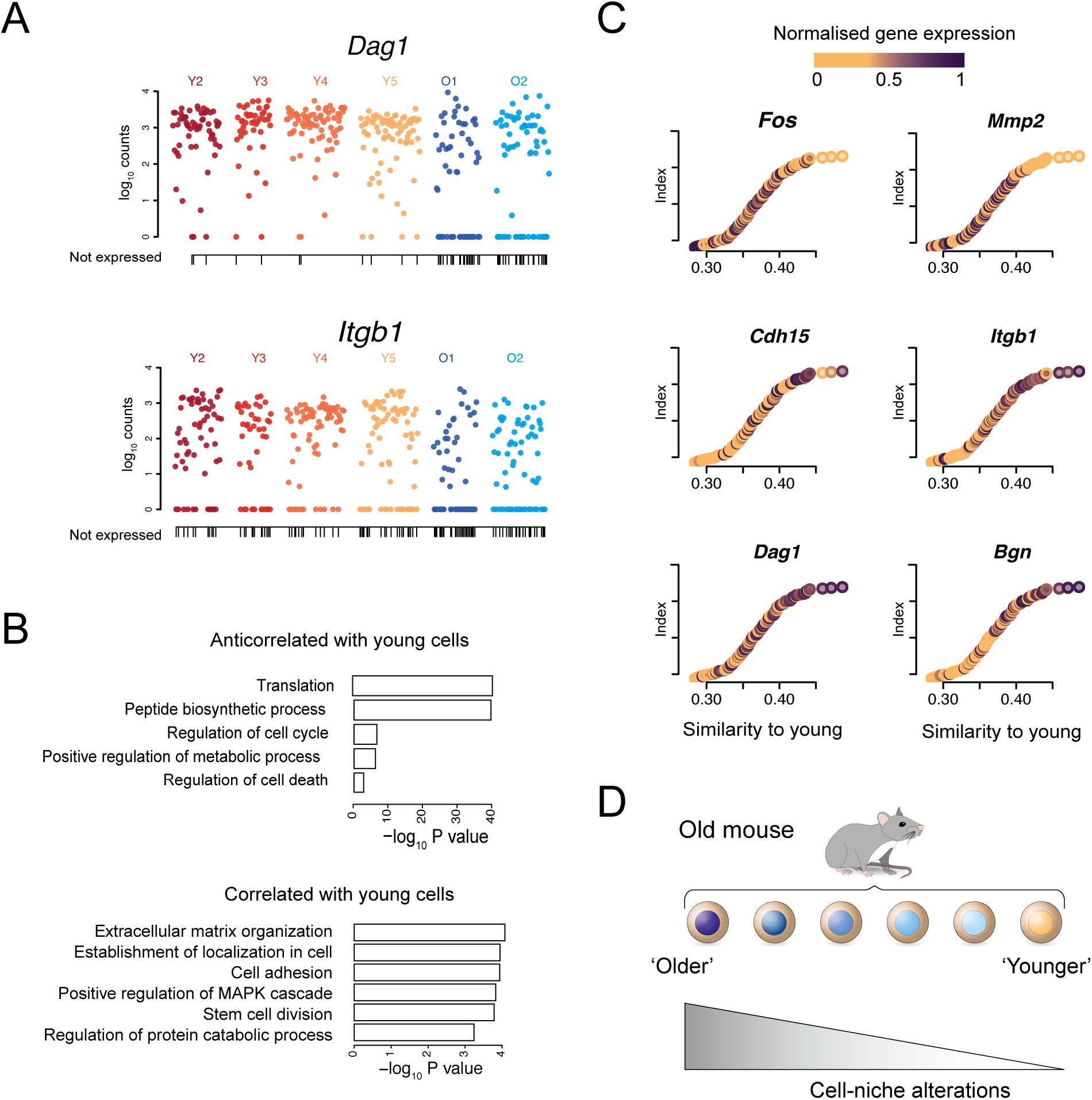
Variability within aged satellite cells and cell-niche interactions. (A) *Dag1* and *Itgb1* expression in young and old cells. Each dot represents a cell. Vertical lines on the x-axis indicate cells that do not express the gene. (B) P-values of the GO terms associated with the top 200 anticorrelated (top) and correlated (bottom) genes with the similarity score to young cells. (C) Similarity between old and young cells. Each dot represents an old cell; on the x-axis cells are ordered according to their similarity (Spearman correlation coefficient) to young cells (young mean expression). Colours indicate the normalized levels of expression of selected genes correlated with the x-axis. (D) Old cells diverging from young cells are likely to have impaired cell-niche interactions.

The observed increase in transcriptional variability with age could reflect the presence of cell subpopulations or be a purely stochastic process. Despite not observing clear substructure (Fig. 1B and Fig. 1D right), we further investigated the origin of this variability by ranking old cells based on their transcriptome-wide similarity to young cells, and performed correlation analyses to identify the genes driving this ranking. Gene ontology analysis indicated that old cells that differed the most from young cells were enriched in processes such as translation and peptide biosynthesis (Fig. 2B top), while old cells that were most similar to young ones were enriched in extracellular matrix-related functions (Fig. 2B bottom). For example, *Fos* and *Mmp2* were preferentially expressed in the most different old cells, while extracellular markers such as *Dag1, Itgb1, Cdh15* or *Bgn* were expressed in the most similar ones (Fig. 2C). These results indicate that cells that have accumulated more differences with age are likely to have impaired cell-niche interactions and are more prone to exit quiescence (Fig. 2D).

For the analysis of DNA methylation patterns, we limited potential biases due to uneven sequencing depth between cells or different number of cells per individual by randomly subsampling 1 million reads from each cell and 35 cells per individual (140 cells in total, 2 million CpG sites on average per cell). Global mean DNA methylation levels were around 50%, as previously reported for muscle stem cells^28^ (Fig. S1C). As with the transcriptomes, we did not observe clear subpopulations in any of the methylome samples (Fig. S2). Overall, CpG islands, promoters and enhancers were hypomethylated; exons, myoblast enhancers (marked by H3K27ac) and shores (flanking region of the CpG islands) were around 30% methylated, while repeats and bodies of active genes (marked by H3K36me3) were highly methylated (Fig. 3A). We found that DNA methylation levels increased slightly with age, as reported for human muscle stem cells^22^, mostly in repeat elements and H3K36me3 regions (Fig. 3B, 3C and 3D).

**Fig. 3.**
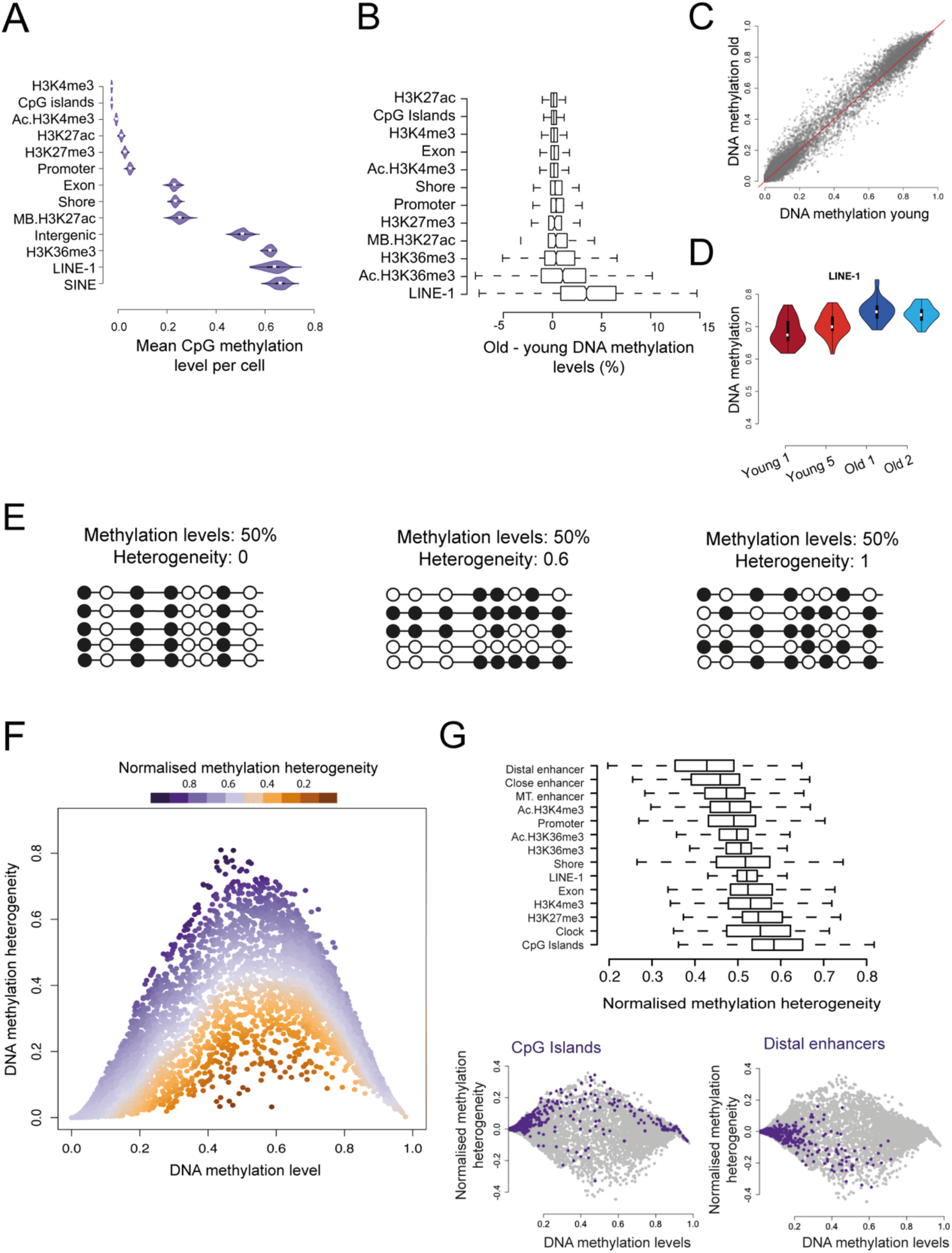
Changes in methylation levels and methylation heterogeneity. (A) Levels of DNA methylation per cell across different genomic regions (Chip-seq data from 2-months-old mice ^23^, Ac: activated satellite cells ^23^, MB: myoblast ^29^). (B) Mean methylation difference between old and young cells across different genomic elements. (C) Genome-wide mean methylation values in old and young cells. Each dot represents a genomic region. (D) Levels of DNA methylation per cell and individual across Line L1 elements. (E) Examples of different distributions of DNA methylation heterogeneity at loci with similar average methylation. Empty circles represent unmethylated CpG sites and filled circles methylated CpG sites. (F) DNA methylation levels and DNA methylation heterogeneity. Each dot represents a genomic region from young or old cells. Colour scale represents the methylation-level normalised measure of DNA methylation heterogeneity. (G) Boxplot showing the normalised DNA methylation heterogeneity across different genomic elements in young cells (top). Normalised methylation heterogeneity and methylation levels across all the different genomic elements (grey) and across CpG Islands (purple) or enhancer regions (purple) in young cells (bottom).

Identical average methylation levels for a given genomic region may reflect different scenarios, from uniform populations to completely random heterogeneous patterns (Fig. 3E). Since we did not observe substructure in our data (Fig. S2) and as stochastic epigenetic drift has been suggested to be a major hallmark of ageing^8^, we computed a score to measure levels of stochastic intrapopulation heterogeneity (Fig. S3, Methods). As expected, our initial measure of heterogeneity depended on average methylation levels (Fig. 3F). Hence, we developed an independent measure of heterogeneity by calculating the distance between the observed heterogeneity for each genomic region and a rolling median (Fig. 3F, Methods). Interestingly, this analysis showed that different genomic contexts displayed different levels of methylation heterogeneity between cells, for example CpG islands were more heterogeneous than enhancers (Fig. 3G).

Global levels of methylation heterogeneity were similar between ages (Fig. S4); we next computed localised Z-score comparisons between young and old to examine changes in specific genomic elements. Notably, methylation of LINE-1 elements became more homogeneous with age whereas regions marked by H3K27me3 became more heterogeneous (Fig. 4A). Specifically, LINE-1 elements also experienced the highest increase in absolute DNA methylation levels, both of which may reflect a coordinated mechanism to prevent deleterious somatic retrotranspositions during ageing. Most of the H3K27me3 regions were associated with genes that are repressed but poised for rapid activation^29^. We hypothesize that this increase in methylation heterogeneity may contribute to an impaired transcriptional response upon activation.

**Fig. 4.**
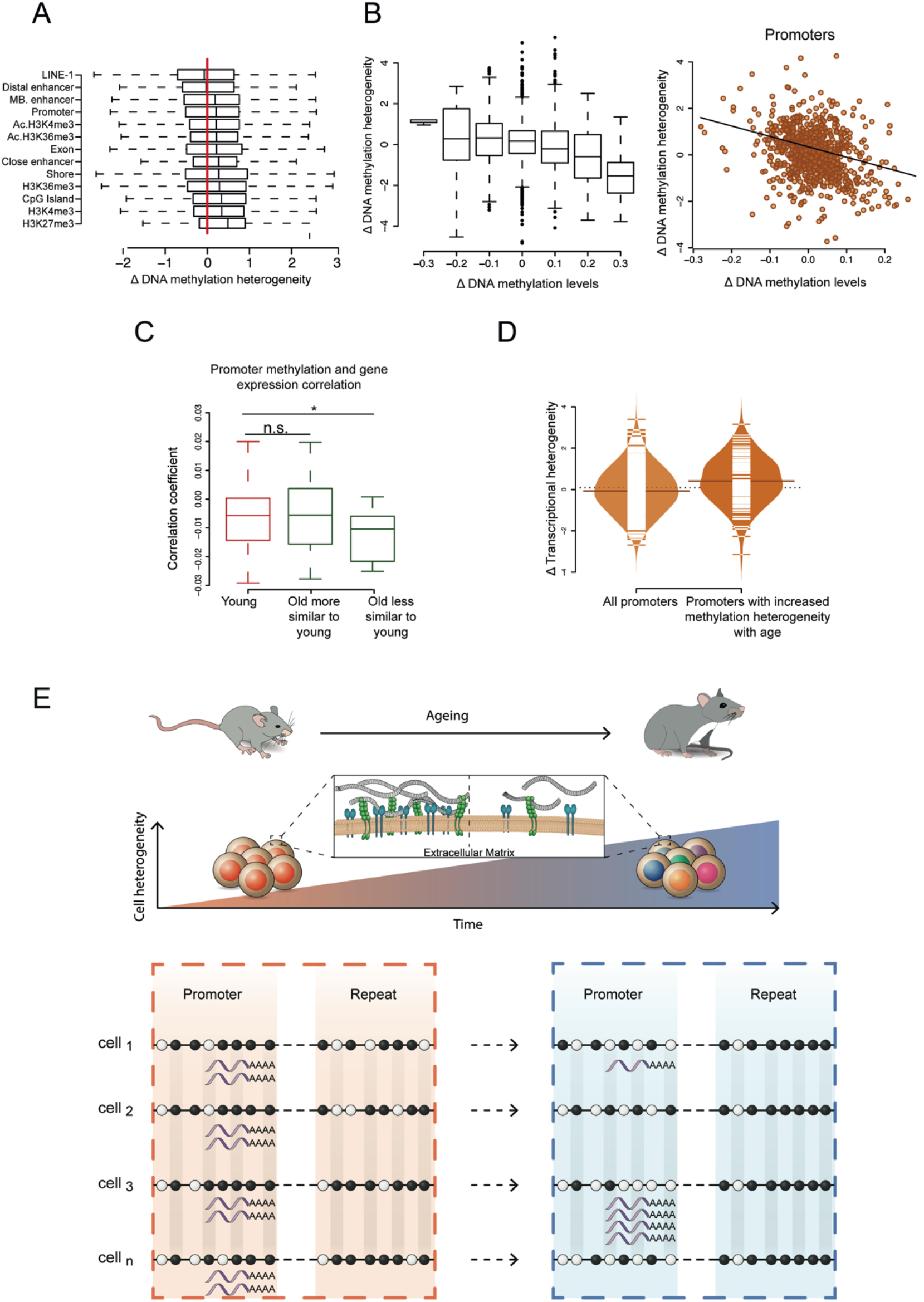
Changes in cell-to-cell methylation heterogeneity during ageing. (A) Normalised methylation heterogeneity changes with age (Δ methylation heterogeneity: old–young) across different genomic features (Ac: activated satellite cells ^23^, MB: myoblast ^29^). (B) Genome-wide normalised methylation heterogeneity difference with ages (Δ methylation heterogeneity: old–young) binned by 0.1 methylation level differences (left). Changes in promoter methylation heterogeneity (y-axis) and methylation levels (x-axis) with age (right). (C) Distribution of Pearson’s correlation coefficients between promoter DNA methylation and gene expression (one association test per cell, number of cells: young = 64, old more similar to young = 30, old less similar to young = 20, * *P* < 0.05). (D) Increase of transcriptional heterogeneity with age across all promoters (n=394) and promoters with increased DNA methylation heterogeneity (Δ methylation heterogeneity > 0.3, n=113) (*P* <0.001). (E) Global increase of transcriptional cell-to-cell variability with age with enhanced heterogeneity in the multiple extracellular matrix related genes (top). Relationship between transcriptional and DNA methylation heterogeneity in aged satellite cells (bottom). Empty circles represent unmethylated CpG sites and filled circles methylated CpG sites. Repeat elements become more homogeneous with age by increasing their methylation levels in a coordinated manner. In contrast, promoter regions become more heterogeneous by randomly loosing DNA methylation and this is coupled with an increase of transcriptional variability of the genes they drive.

Interestingly, we observed a negative correlation between changes in methylation levels and changes in methylation heterogeneity (Promoters: Pearson’s coefficient= −0.35, *P* < 2.2e-16, Fig. 4B). Regions becoming more homogeneous showed an increase in methylation, suggesting that *de novo* methylation enzymes (Dnmt3a,b) are recruited to specific sites and add methylation in a coordinated manner between cells. In contrast, regions becoming more heterogeneous showed a decrease in their methylation levels. Despite the low proliferative history of these cells, this pattern could reflect errors in DNA methylation maintenance during DNA replication, or an active demethylation mechanism via TET enzymes (Fig. S5).

Epigenetic changes may contribute to the age-associated pattern of transcriptional heterogeneity. To explore this possibility, we analysed the association between promoter DNA methylation and gene expression. We calculated a correlation coefficient for each cell and confirmed the expected negative correlation for methylation and transcription (Fig. 4C). Interestingly, old cells that were most transcriptionally different from young cells showed lower levels of correlation (Mann-Whitney-Wilcoxon test; *P* < 0.05, Fig. 4C). Furthermore, we calculated changes in transcriptional variability between young and old cells (see Methods) and observed that promoters with increased methylation heterogeneity tended to have increased transcriptional heterogeneity (Mann-Whitney-Wilcoxon test; *P* <0.001) (Fig. 4D). It appears therefore that deterioration of transcriptional coherence during ageing is associated with increased promoter methylation heterogeneity and with decreased connectivity between the epigenome and the transcriptome.

In summary, we report transcriptional and epigenetic signatures associated with ageing in a deeply quiescent population of muscle stem cells. Previous studies have investigated transcriptional heterogeneity changes with age in mixed cell populations^4^ which are affected by differences in cellular composition, such as an increase in senescent cells^4^. In contrast, our study is focused on a specific population of cells in which known stemness, activation and senescent markers were not affected by ageing. Even in this restricted population, we observe a global increase of uncoordinated transcriptional variability with age, indicating an intrinsic mechanism of cellular ageing. Interestingly, mouse muscle stem cells were shown to maintain clonal diversity during homeostatic ageing by lineage-tracing^18^, however, our study uncovers a dramatic underlying molecular heterogeneity in these stem cells that extends beyond maintenance of clonal homogeneity. We also observe that cells that have acquired more differences with age showed alterations in multiple extracellular matrix related genes potentially affecting cell-niche interactions.

Elevated transcriptional variability with age has been reported in several studies^2–4^, however the underlying causes remain largely unknown. The accumulation of somatic mutations only partially accounts for the increased cell-to-cell transcriptional variability^4^, suggesting that epigenetic mechanisms might be a contributing factor^5^. In this study, by applying for the first time a combined single cell method for DNA methylation and the transcriptome, we show that epigenetic drift, or the uncoordinated accumulation of methylation changes in promoters, contributes to the increased transcriptional variability with age (Fig. 4E). Due to the deep quiescent state of the homeostatic cells chosen for study, our data highlight the possibility that the observed epigenetic patterns could be independent of extensive cell proliferation. We propose that this variability is detrimental due to uncoordinated transcription, thereby affecting the ability of stem cells to maintain quiescence or activate coherently upon injury. Future studies of different stem cell populations integrating multiple layers of molecular information will be highly informative for a more complete understanding of the underlying molecular mechanisms of ageing and age-related diseases.

## Methods

### Mice

Animals were handled according to national and European Community guidelines, and an ethics committee of the Institut Pasteur (CETEA) in France approved protocols. Young (2 months-old) and old (24 months-old) *Tg:Pax7-nGFP*^21^ mice were used in this study.

### Isolation of satellite cells

Mice were sacrificed by cervical dislocation. *Tibialis anterior* muscles were dissected and placed into cold DMEM (ThermoFisher, 31966). Muscles were then chopped and put into a 15 ml Falcon tube containing 10 ml of DMEM, 0.08% collagenase D (Sigma, 11 088 882 001), 0.1% trypsin (ThermoFisher, 15090), 10 *μ*g/ml DNaseI (Sigma, 11284932) at 37°C under gentle agitation for 25 min. Digests were allowed to stand for 5 min at room temperature and the supernatants were collected on 5 ml of foetal bovine serum (FBS; Gibco) on ice. The digestion was repeated 3 times until complete digestion of the muscle. The supernatants were filtered through a 70-*μ*m cell strainer (Miltenyi, 130-098-462). Cells were spun for 15 min at 515g at 4°C and the pellets were resuspended in 1 ml freezing medium (10% DMSO (Sigma, D2438) in foetal calf serum (FCS, Invitrogen)) for long term storage in liquid nitrogen.

Before isolation by FACS, samples were thawed in 50 ml of cold DMEM, spun for 15 min at 515g at 4°C. Pellets were resuspended in 300 *μ*l of DMEM 2% FCS 1 *μ*g/mL propidium iodide (Calbiochem, 537060) and filtered through a 40-*μ*m cell strainer (BD Falcon, 352235). Viable cells were isolated based on size, granulosity and GFP expression level (top 10% nGFP^Hi^ cells, Fig. S6) using a MoFlo Astrios cell sorter (Beckmann Coulter).

Single cells were collected in 2.5 *μ*L cold RLT Plus buffer (Qiagen, 1053393) containing 1U/*μ*L RNAse inhibitor (Ambion, AM2694) in 96 well-plates (LoBind Eppendorf, 0030129504), flash-frozen on dry ice and stored at −80°C.

### Library preparation and data alignment

We prepared scM&T-seq libraries^10^ by isolating mRNA on magnetic beads and separating from the single-cell lysate as described^30^ prior to reverse transcription and amplification using Smartseq2^31^ but with 25 PCR cycles. We then processed the lysate containing genomic DNA according to the published single-cell bisulfite sequencing protocol^32^. Single-cell RNA-seq libraries were aligned using HiSat2 with options–sp 1000,1000–no-mixed–no-discordant^33^. Single-cell bisulfite libraries were processed using Bismark^34^ as described^10^. Mapped RNA-seq data were quantitated using the RNA-seq quantitation pipeline in Seqmonk software (www.bioinformatics.babraham.ac.uk/projects/seqmonk/).

### Quality control RNA-seq

Cells expressing fewer than 1,000 genes or less than 10^5^ mapped reads allocated to nuclear genes were removed in quality control (Fig. S7). These cells were also verified to have less than 10% of mapped on mitochondrial genes. Out of the 768 cells that were captured across the experiment, 377 passed our quality and filtering criteria (Table S2).

### Data analysis RNA-seq

Gene expression levels were estimated in terms of reads per million of mapped reads to the transcriptome. A score of variability per gene (named distance to the median) was calculated by fitting the squared coefficient of variation as a function of the mean normalized counts and then calculating the distance to a rolling average (window size=100) (Fig. S8)^24^. We included only genes with an average normalized read count of at least 10. The top 1000 most variable genes of the entire data set were used to perform principal component analyses (as log_2-_transformed and median-cantered values) (Fig. 1B, Table S3). Single cell differential expression (SCDE) was used to calculate differential expression analysis between young and old cells (Table S1)^35^.

Cell-to-cell correlation analyses were performed using the top 500 most variable genes within each individual and using Spearman’s correlation as the measure of similarity between cells (Fig. 1D). Distance to the median of the top 500 most variable genes within each individual was computed for Fig. 1E, similar results are observed when restricting the analysis to genes that are expressed in all the individuals (average normalized read count of at least 10) and different numbers of genes (Fig. S9).

An average young reference transcriptome was computed by calculating the mean of log transformed expression values for each gene across cells from young individuals. We then performed Spearman’s correlation analyses to assess the similarity between each cell from old samples and the young transcriptome. Spearman’s correlation analyses were then also used to find gene expression patterns associated with this genome-wide similarity score. Genes expressed in fewer than five cells were excluded from the analysis. The top 200 correlated and anticorrelated genes (Table S4) were used for GO enrichment analysis^36^.

### DNA-methylome

We discarded cells that had less than 1 million paired-end alignments or less than 500,000 CpG sites covered (Fig. S1). To avoid biases that might occur due to different sequencing depths or number of cells between individuals, we down-sampled the data to 1 million reads for each cell and randomly selected 35 cells from each individual (2 young and 2 old). ChIP-seq datasets for H3K4me3, H3K27me3, H3K36me3 in satellite cells and H3K27ac in myoblast were obtained from existing studies^23,29^. Bowtie2 and MACS2 were used for mapping and peak calling respectively.

### DNA methylation heterogeneity

We developed a heterogeneity score based on Hamming distances and Shannon entropy between cell pairs from the same sample. This value captures the properties we desire: i) ability to detect cell-to-cell stochastic heterogeneity ii) not affected by population substructure iii) not biased by missing values. Precisely, let *r* be a matrix with methylation values of cells for a particular gene, each row corresponding to a cell and each column corresponding to a CpG site, and *w* be the weight corresponding to the number of covered CpGs within each pairs of cells. For each pair of cells (*c*), we then computed the Hamming distance (*D*) and the Shannon entropy score of the pairs (*S*) considering sites with coverage in both cells. Then weighted heterogeneity score of the regions is:

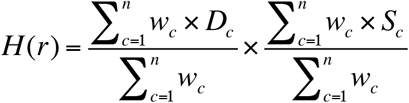

Here *D*_*c*_ is the normalised Hamming distance of a given a pair of cells, which measures the number of bits that are different in two binary sets:

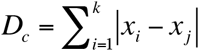

*S*_*c*_ is the joint Shannon Entropy between a pair of cells which measures the complexity of the pattern:

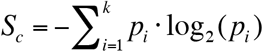

Here *p* is the frequency of pairs of methylation values.

We validated our approach by applying the method in simulated data with increasing levels of methylation heterogeneity (Fig. S3). We also observed that our algorithm is highly robust to missing data (Fig. S3).

We applied this method across multiple genomic regions for each individual independently and then computed the average of young and old samples. Pairwise comparisons with fewer than 4 CpG sites were not considered in the analysis. Furthermore, to avoid misinterpretations because of poor coverage depth we excluded regions with: i) less than 20CpG sites, ii) less than an average of 2 CpG sites covered per cell, iii) less than 100 cell-to-cell pairwise comparisons. We also excluded regions with high coverage differences between ages (more than an average of 10 CpG sites or more than 200 cell-to-cell pairwise comparisons). A total of 63,823 genomic regions were used in the analysis (average window size= 2,267 bp).

Coverage-weighted cell methylation values were used to calculate the mean methylation levels of each region. A normalised measure of DNA methylation heterogeneity was calculated for each region (from young or old samples) by fitting the score of heterogeneity as a function of the mean methylation levels and then calculating the distance to a rolling median of 1,000 observations (Fig. 3F). Regions with less than 0.05 or more than 0.9 mean methylation levels were excluded from the analysis.

Differences between young and old DNA methylation heterogeneity values were Z-score normalised using a sliding window of 100 observations ordered by the mean value of young and old (Fig. S10 and Table S5). Same approach was used to calculate differences between young and old transcriptional heterogeneity (mean distance to the median) (Fig S9 and Table S5).

### Data availability

Sequencing data have been deposited in GEO with the accession: GSE121364

### Software

Custom software is available upon request.

## Acknowledgments

We would like to thank the Flow Cytometry Platform of the Center for Technological Resources and Research (Institut Pasteur) and the Wellcome Trust Sanger Institute sequencing facility for assistance with Illumina sequencing. This project was supported by grants from Institut Pasteur, Agence Nationale de la Recherche (Laboratoire d’Excellence Revive, Investissement d’Avenir; ANR-10-LABX-73), Association Française contre les Myopathies (21857), CNRS, the European Research Council (Advanced Research Grant 332893), and the Biotechnology and Biological Sciences Research Council (BBSRC, CBBS/E/B/000C0425).

## Author contributions

I.H.H., B.E., T.S., S.T. and W.R. proposed the concept and designed the experiments. B.E and P.H.C. performed FACS; T.S. and S.C. performed library preparation and sequencing. I.H.H. developed the analysis methodologies and analysed the experiments with advice from SA. I.H.H., B.E., S.T. and W.R. wrote the paper. All authors read and agreed on the manuscript.

## Materials & Correspondence

Correspondence and material requests should be addressed to.

---

**Fig. S1.**
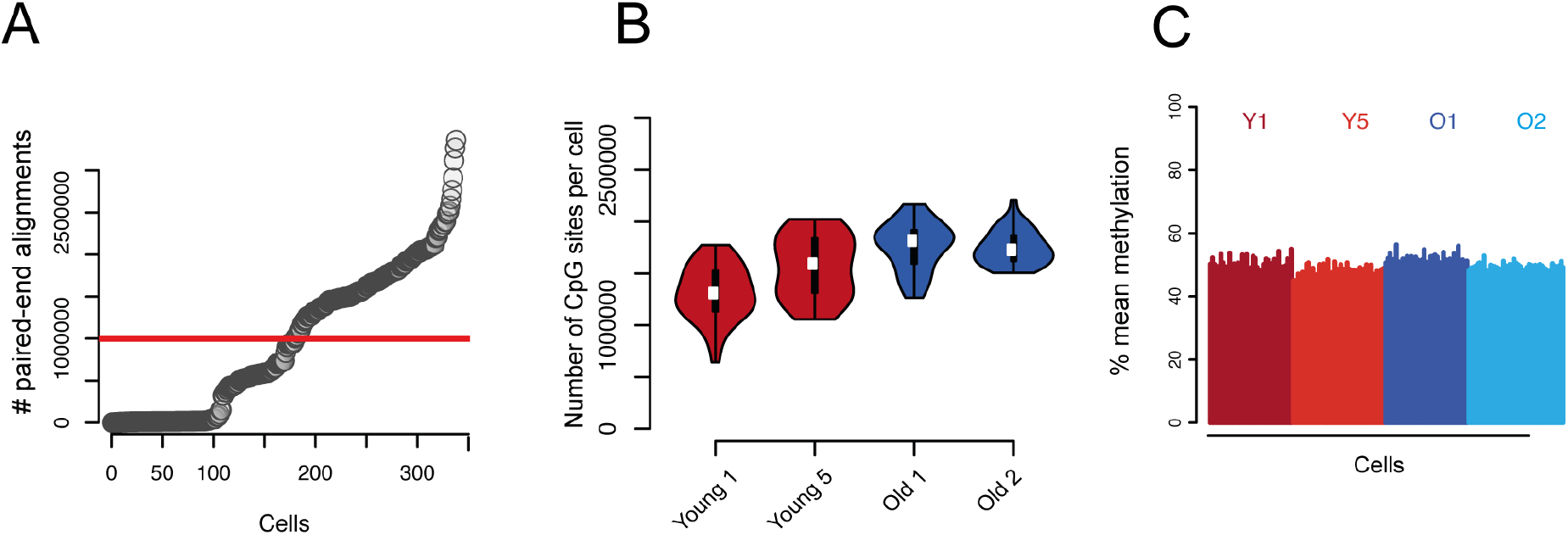
Quality control of single-cell DNA methylation data. (A) Number of pair-end alignments per cell. Cells below the threshold were excluded from the study. (B) Number of CpG sites per cell and individual. (C) Mean methylation per cell showing no global differences between ages.

**Fig. S2.**
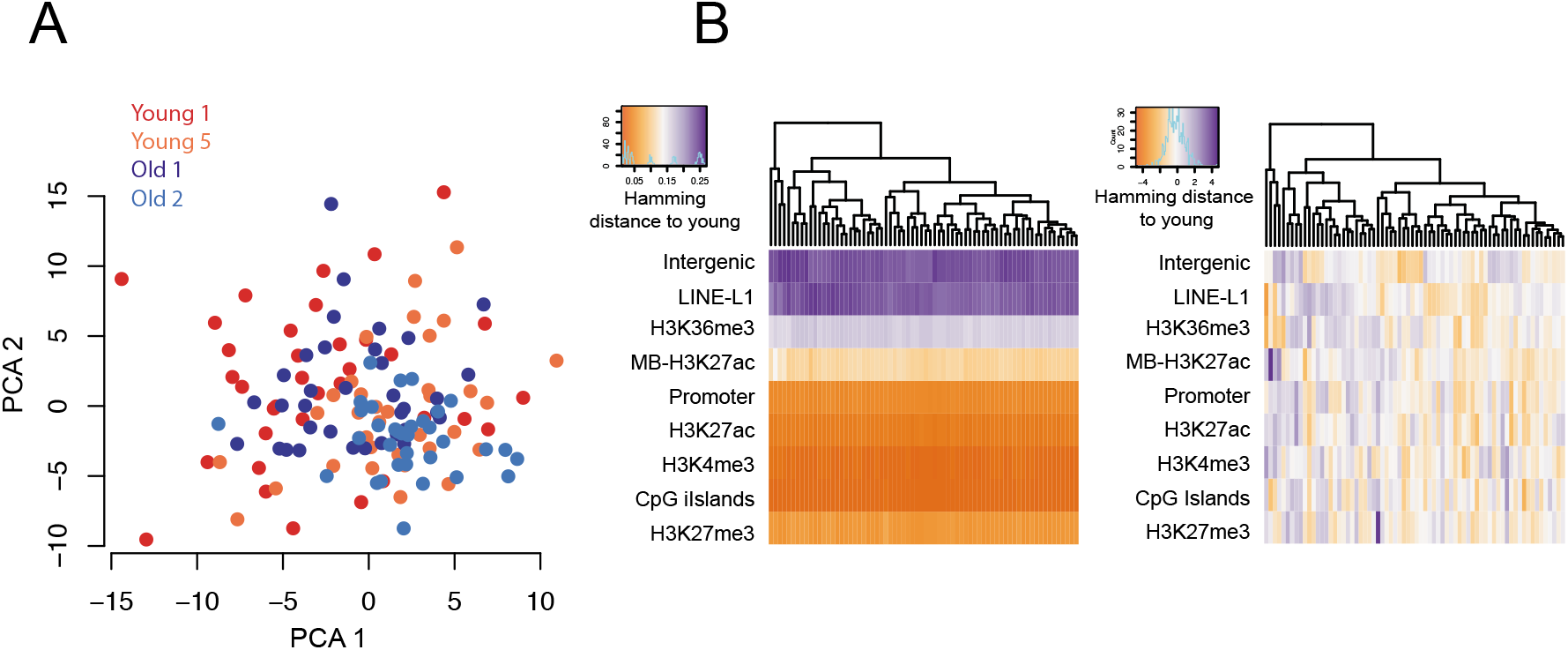
Cell clustering based on DNA methylation data. (A) PCA on gene body methylation showing no clear differences between ages. (B) Heatmap showing Hamming distances between the average methylation from young cells and individual old cells (columns) across different genomic context (rows) (left). Same measure normalised by genomic context (right) showing no cellular substructure.

**Fig. S3.**
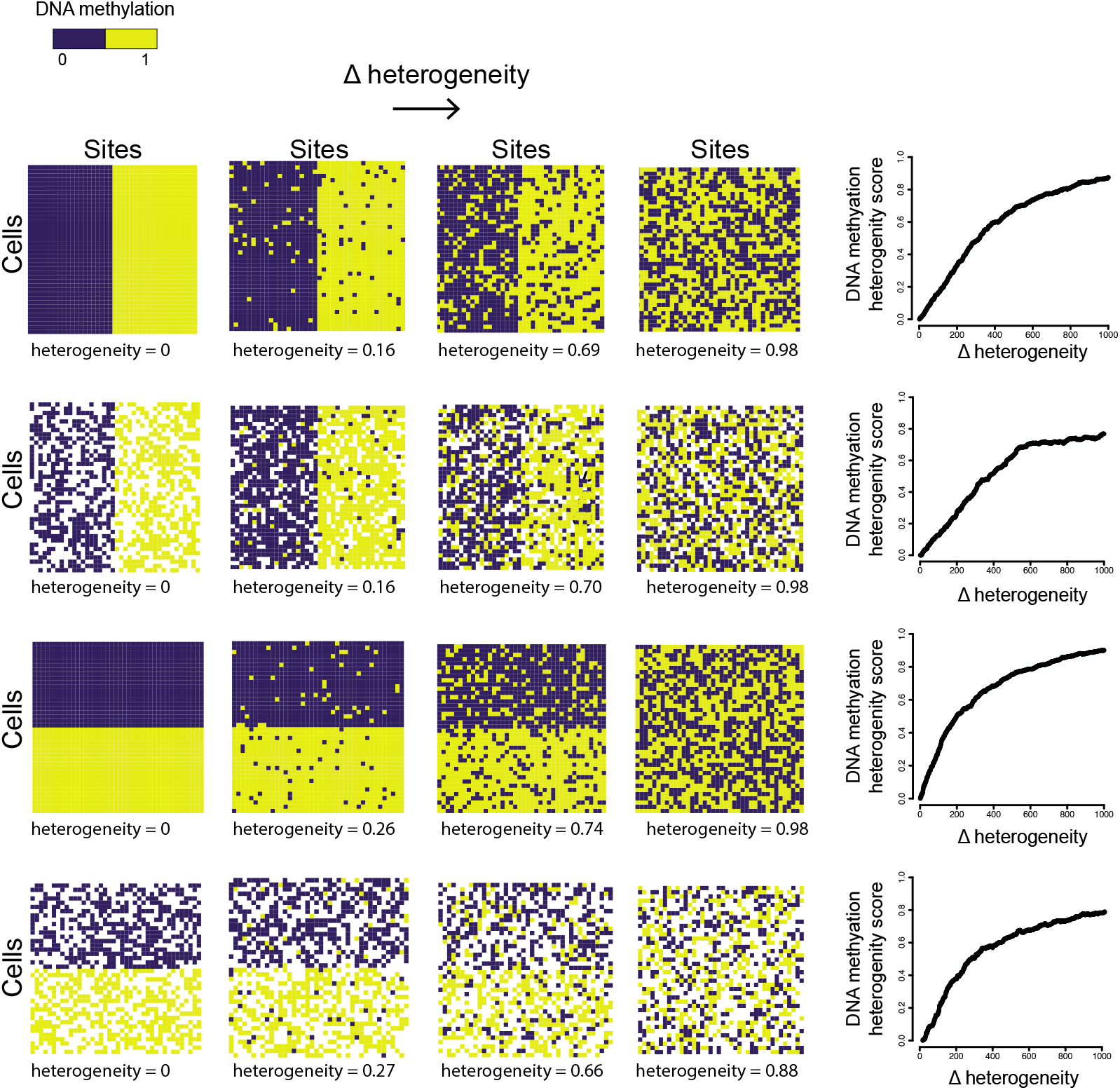
DNA methylation heterogeneity on simulated data. Four cases with different population substructure and missing values tested with simulated data of increasing heterogeneity. Missing values are represented in white.

**Fig. S4.**
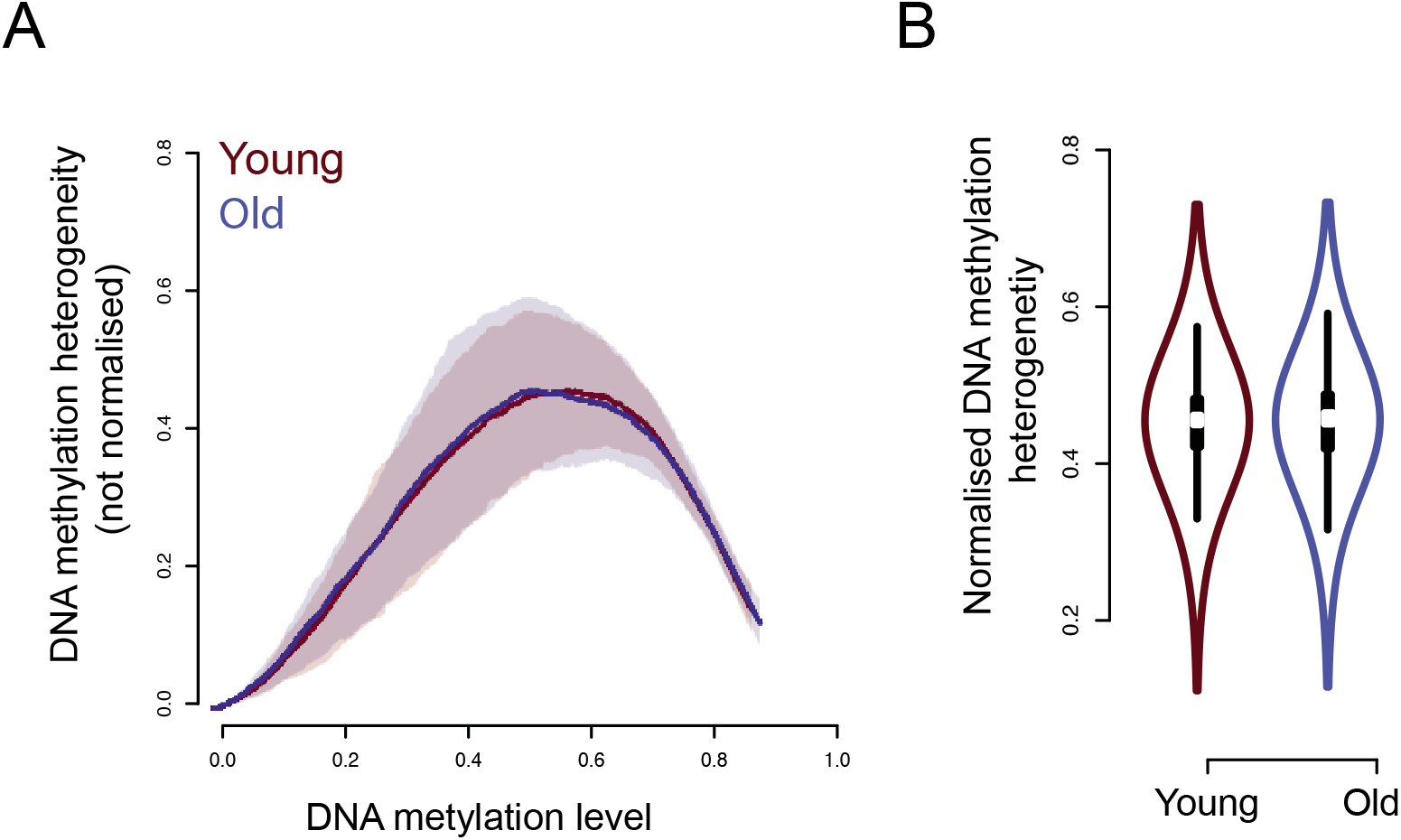
Global levels of DNA methylation heterogeneity between ages. (A) DNA methylation levels and methylation heterogeneity in young and old cells. (B) Normalised DNA methylation heterogeneity in young and old cells.

**Fig. S5.**
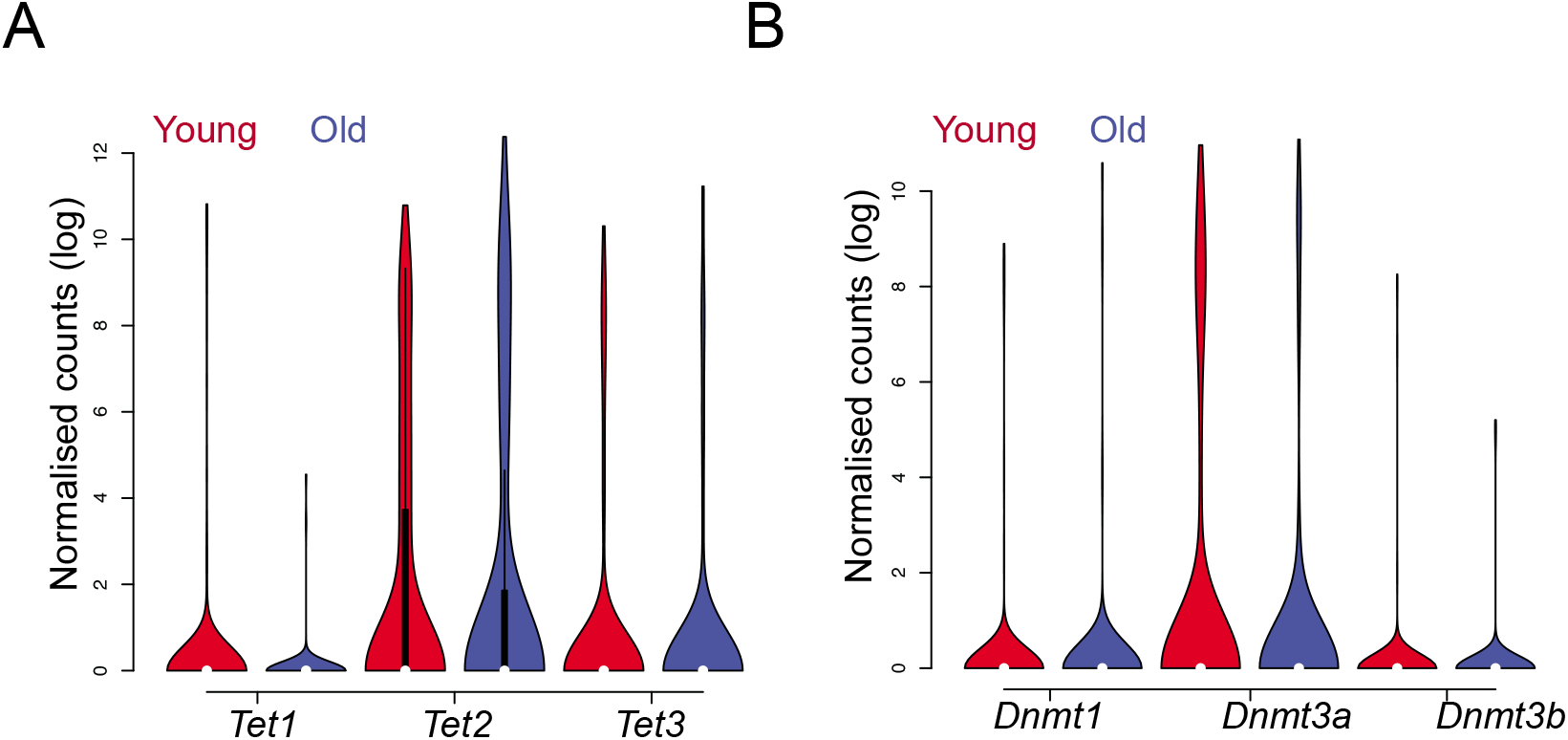
Expression levels of the DNA methylation enzymes. (A) Expression levels of the enzymes for active demethylation in young and old samples. (B) Expression levels of the DNA methylation enzymes in young and old samples.

**Fig. S6.**
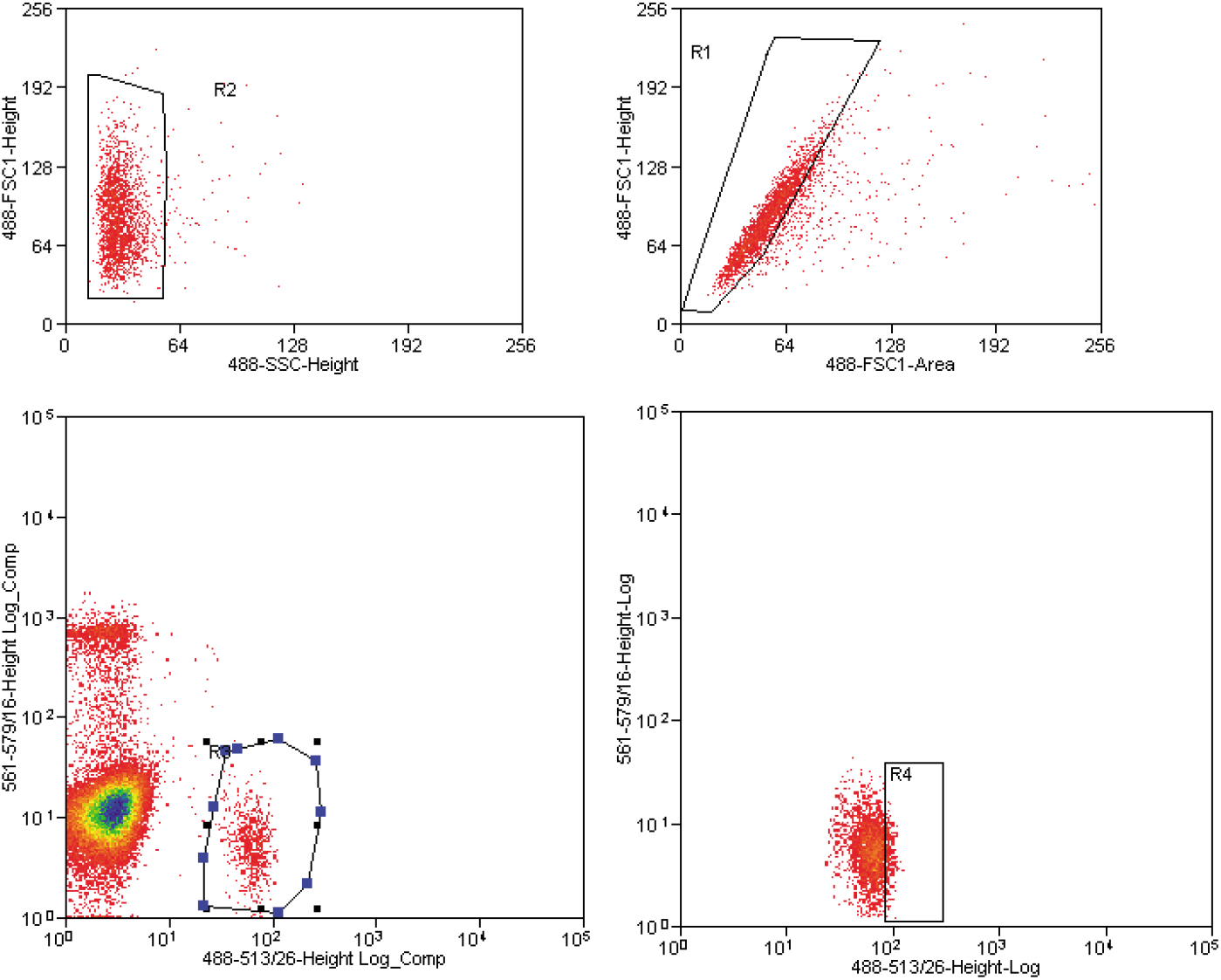
Isolation of single satellite cells by FACS. Satellite cells were isolated by FACS by gating first on size and granulosity (R2 gate), excluding doublets (R1 gate) and gating on the GFP^+^/PI^-^ population (R3 gate). Pax7-nGFP^Hi^ cells (top 10% highest nGFP-expressing cells, R4 gate) were sorted as single cells.

**Fig. S7.**
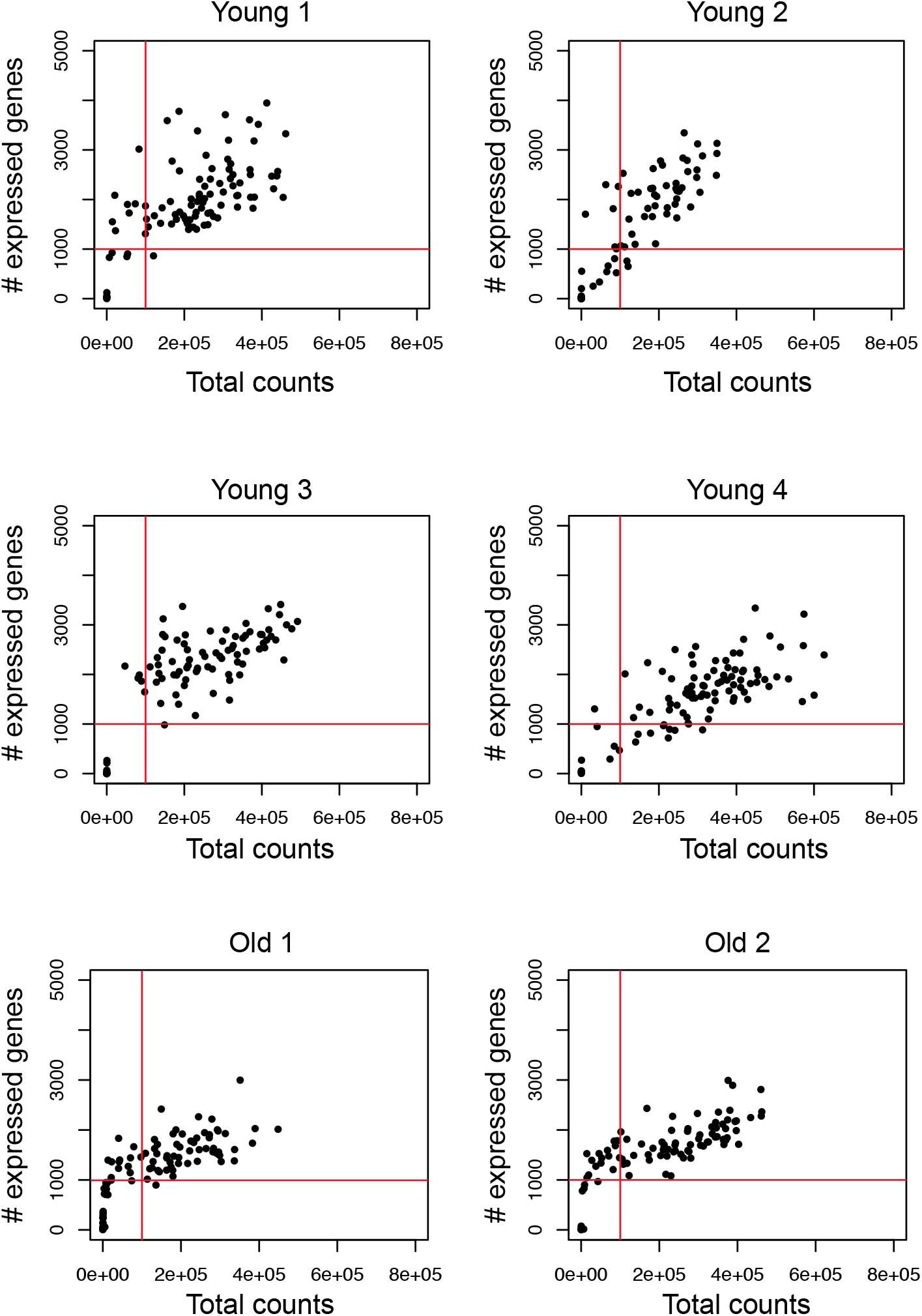
Quality control of single-cell RNA-seq data. Plot representing number of genes and total expression counts expressed in each cell per individual. Cells above highlighted threshold (1000 genes, 10^5^ counts) were included in the study.

**Fig. S8.**
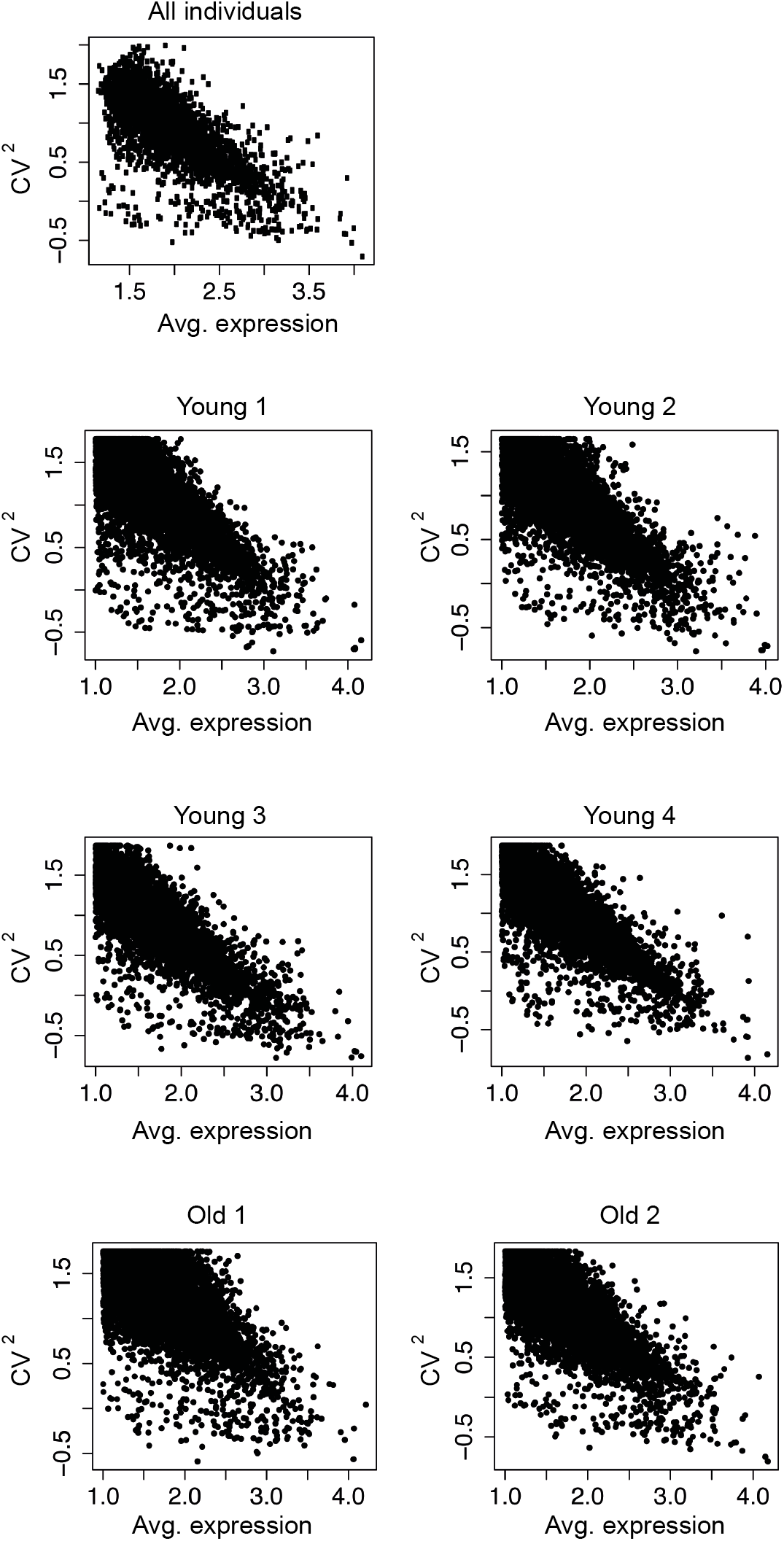
Transcriptional variability. Gene variability: squared coefficients of variation are plotted against the means of normalized read counts for gene using data from all individuals (top) or each individual separately.

**Fig. S9.**
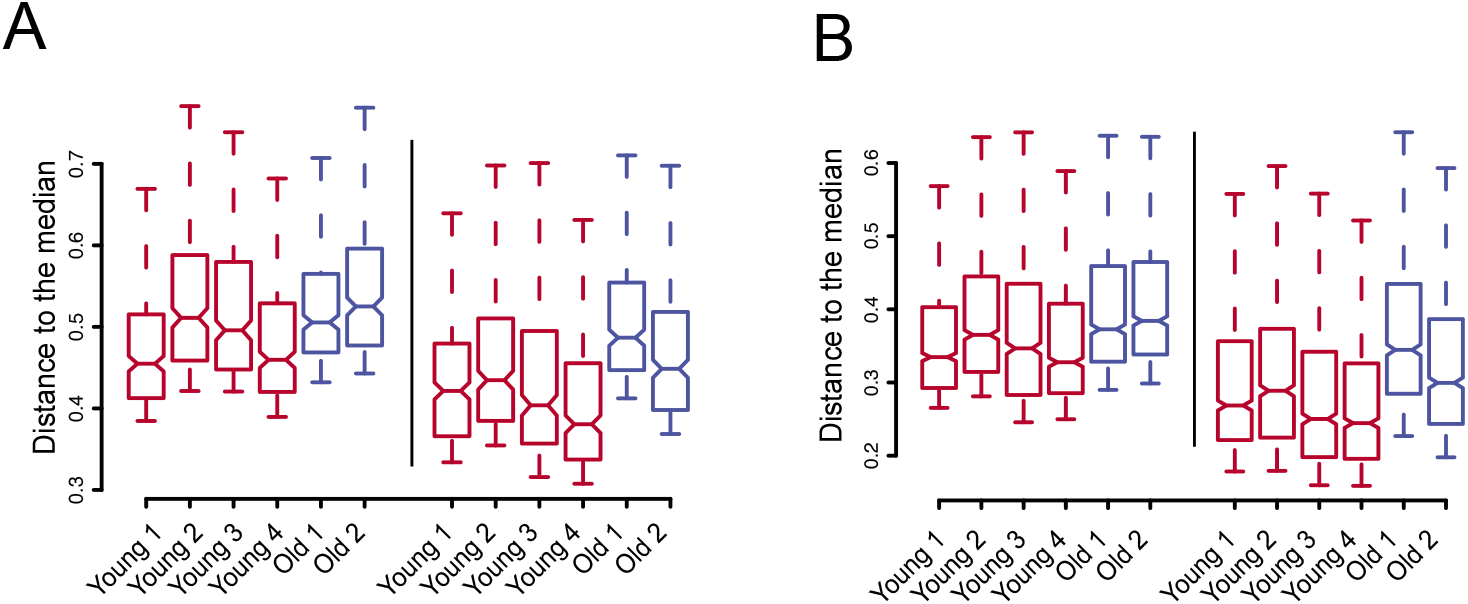
Transcriptional variability: distance to the median. Distance to the median of the top 300 (A) and 1000 (B) most variable genes among all genes (left) and among the 5,127 common genes expressed in the six individuals (right).

**Fig. S10.**
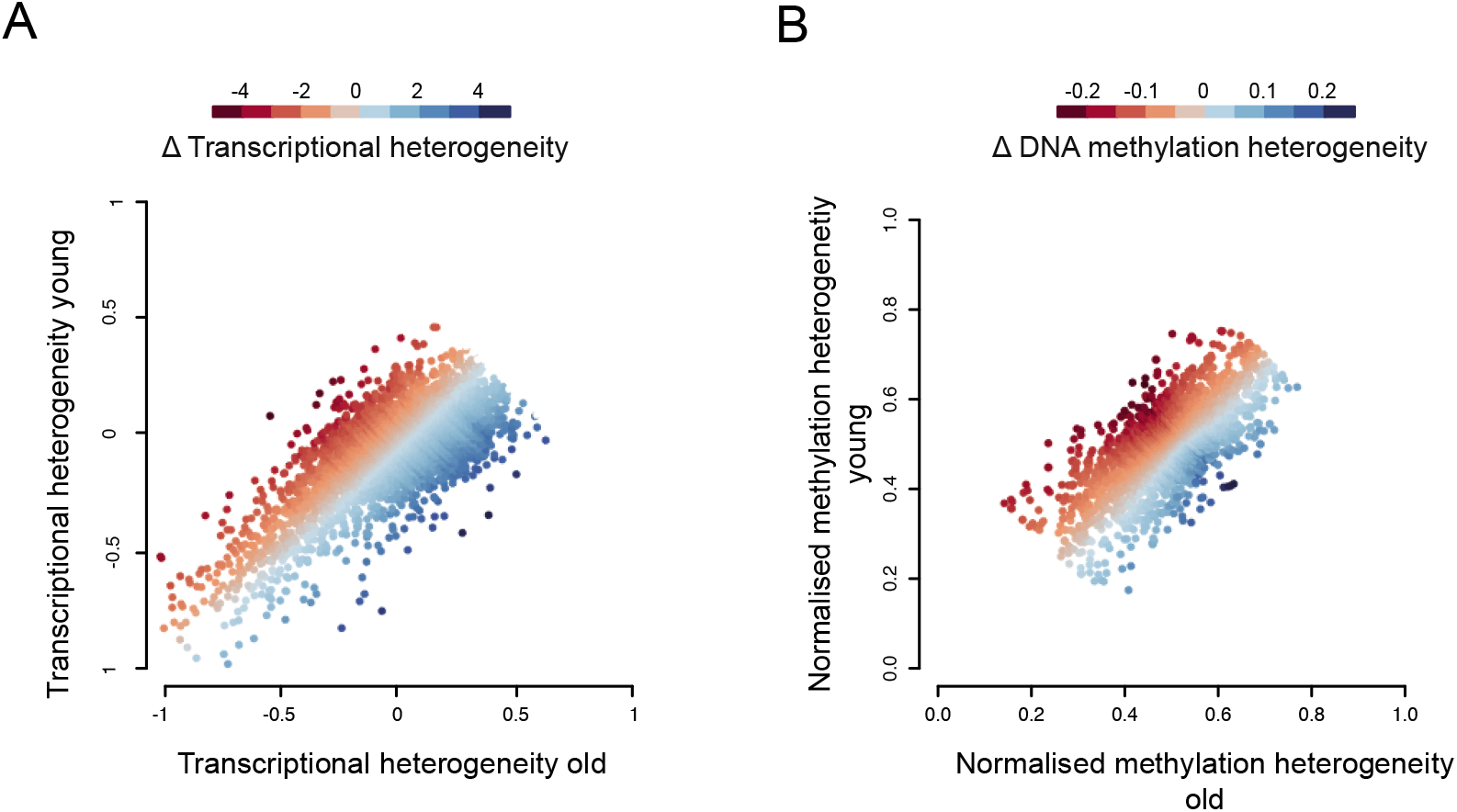
Changes in transcriptional and DNA methylation heterogeneity with age. (A) Differences in transcriptional heterogeneity measures where Z-score normalised using a sliding window of 100 observations (color code). Transcriptional heterogeneity represents the mean distance to the median for every gene from young (y-axis) and old (x-axis) individuals. (B) Differences in DNA methylation heterogeneity measures where Z-score normalised using a sliding window of 100 observations (color code). DNA methylation heterogeneity represents the normalised measure of methylation heterogeneity from young (y-axis) and old (x-axis) individuals.

**Table S1.** Differentially expressed genes between cells from young and old mice.

**Table S2.** scM&T quality control.

| Sequencing label | Individual | RNA |  | DNA |  |
| --- | --- | --- | --- | --- | --- |
|  |  | Total cells | After QC | Total cells | After QC |
| Y2 | Young 1 | 96 | 0 | 78 | 35 |
| Y8 | Young 2 | 96 | 75 | 86 | 35 |
| Y5 | Young 3 | 96 | 44 | NA | NA |
| Y7 | Young 4 | 96 | 74 | 72 | 35 |
| Y4 | Young 5 | 96 | 60 | NA | NA |
| O1 | Old 1 | 96 | 56 | 80 | 35 |
| O5 | Old 2 | 96 | 68 | 90 | 35 |
| O8 | Old 3 | 96 | 5 | 8 | 0 |

**Table S3.** Top 1000 most variable genes across the entire data set.

**Table S4.** Top 200 genes correlated and anticorrelated with the similarity score to the young reference transcriptome.

**Table S5.** Increase of transcriptional and methylation heterogeneity with age in promoter regions (Δ: Old-young).

